# Temporal Dynamics of Neural Processing of Facial Expressions and Emotions

**DOI:** 10.1101/2021.05.12.443280

**Authors:** Sanjeev Nara, Dheeraj Rathee, Nicola Molinaro, Naomi Du Bois, Braj Bhushan, Girijesh Prasad

## Abstract

Emotion processing has been a focus of research in psychology and neuroscience for some decades. While the evoked neural markers in human brain activations in response to different emotions have been reported, the temporal dynamics of emotion processing has received less attention. Differences in processing speeds, that depend on emotion type, have not been determined. Furthermore, behavioral studies have found that the right side of the human face expresses emotions more accurately than the left side. Therefore, accounting for both the content of the emotion and the visual angle of presentation from the perspective of the viewer, here we have investigated variability in the discrimination of happy and sad faces when the visual angle of presentation was Positive (right side of the face) compared to Negative (left side of the face). Furthermore, the temporal dynamics involved in processing happy and sad emotions have been compared. Regardless of visual angle, happy emotions were processed faster than sad emotions. However, the evoked response to sad emotions significantly increased in amplitude compared to that for happy emotions, when faces were presented at Positive visual angles only. Source reconstruction from sensor-level ERFs show localized activities in ventral and dorsal stream, including fusiform gyrus, lingual gyrus, putamen and Pre and Post central gyrus. Multivariate pattern analysis (MVPA) confirmed these findings – demonstrating successful decoding of happy and sad emotions only occurred when the facial expression was viewed from a positive visual angle, and that happy emotions were processed faster than sad emotions.

## Introduction

Emotion expression recognition is an important skill for human survival and social interactions, providing important information regarding potential threat while interacting with humans and other species. Ekman and colleagues have proposed six basic emotions (Ekman et al., 1987). It is evident from behavioral studies that there are hemispheric asymmetries inherent in facial expression processing.

Researchers have discussed the right hemisphere dominance for processing of facial expressions of emotion. While the right-hemisphere hypothesis has proposed right hemisphere dominance for all emotions (Heilman and Bowers, 1990; Yecker et al., 1999), the valence hypothesis suggests that the right and left hemispheres are dominant for negative and positive emotions (Silberman and Weingartner, 1986), respectively. Due to contralateral control, several behavioral studies (Mandal and Ambady, 2004; Bhushan, 2006) have reported hemifacial bias. These studies demonstrate that the left side of the face distinctly expresses emotions, especially negative emotions. However, Bryden (Bryden M. P., 1982) has argued that the hemispheric dominance pertains to expression and experience of emotion and not perception of the emotion. Therefore, irrespective of valence, the right hemisphere is dominant for the perception of emotion (Davidson, 1984). It merits mentioning that the research to date has relied on responses to facial expression presented using frontal views of the face only (referred to here as a 0° visual angle). It is an important yet neglected consideration that poses a serious limitation to such generalization, as in real life we come across many situations when the visual angle is either Negative (left faces) or Positive (right face).

The Indian Affective Picture Database (IAPD) (Sharma and Bhushan, 2019) and the Karolinska Directed Emotional Faces (KDEF) (Lundqvist et al., 1998) are the only databases that contain pictures of seven emotional expressions taken from four different angles, i.e., −90° (full left profile), −45° (half left profile), +45° (half right profile) and +90° (full right profile). However, the left (−90° and −45°) and right (+45° and +90°) profiles of these databases have yet to be examined for the left/right facial asymmetry that has been so well documented in behavioral studies and neurosciences. Therefore, in this study we aim to investigate the left/right facial asymmetry for facial expressions of emotions spanning from full left to full right profile view, using a state-of-the-art neuroimaging technique – Magnetoencephalography (MEG). Applying fractal dimension to IAPD images, Bhushan and Munshi (Bhushan and Munshi, 2021) found that recognition of happy emotion does not get affected by visual angle. If visual angle does not affect recognition of happiness, why is the opposite of happiness (sadness) affected by it? MEG can unravel the hemispheric asymmetry along with localizing the site of such processing.

Emotion processing has been studied in detail in the last few decades. Electroencephalography (EEG) neuroimaging studies have revealed evoked potential responses (ERPs) that have been used extensively to understand emotion processing, with three main ERP stages proposed by Ding and colleagues (Ding et al., 2017); a first stage for automatic but coarse processing (N100 and P100), a second stage for distinguishing emotional and neutral facial expressions (N170 and VPP), and a third stage for distinguishing various emotional facial expressions (N300 and P300). There are studies both in favor (Hung et al., 2010) and against (Leppänen et al., 2007) these components, which have been replicated in different geographical locations.

The present study aimed to examine both facial asymmetry and emotion processing mechanisms:

i. to investigate the neurophysiological evidence for hemispheric and hemifacial asymmetry in the processing of facial expressions of emotions, i.e., whether the left or right side of the human face provides equal or different information for emotion processing, and which hemisphere is activated during this processing.
ii. to investigate the time frame needed for the processing of happy and sad expressions of emotions.

Given the temporal resolution required, MEG was used for data collection, while analyses combined traditional ERF, and modern state of the art, analysis techniques like multivariate pattern analysis (MVPA) (Carlson et al., 2019).

## Methods

### Participants

Thirteen participants (*n* = 2, female) took part in the present study (age range: 21 – 38 years; *M* = 27.30; *SD* = 6.61). The ethical committee and scientific committee of Ulster University approved the experiment (following the principles of the Declaration of Helsinki). Participants gave written informed consent and were financially compensated. The participants were recruited from the students enrolled in different courses at Ulster university, UK. Participants were free from any neurological and psychological disorders and had normal or corrected to normal vision.

### Experimental design

The participants were presented with images eliciting happy and sad emotions at four different visual angles (i.e., - 45^0^, -90^0^, +45^0^ and +90^0^) selected from the Indian Affective Picture Database (IAPD) (Sharma and Bhushan, 2019). IAPD comprises 140 colored pictures modelling facial expressions of emotions at five different visual angles (−90, -45, +45, and +90). Each emotion was modelled by four different actors (two male and two female). Figure 1 shows one such actor modelling the facial expressions assigned to both the emotions of happy and sad, presented at all four visual angles. Written consent was taken from the inventors of IAPD database to share the facial expression and images used in this experiment. Two blocks were presented to the participants consisting of 32 trials per emotion and angle leading to a total of 256 trials per participant. Every trial began with a red color cross fixation for a randomly assigned duration of 2000 - 3000 ms, then a face image appeared for 1000 ms. The participants were asked to report the emotion (happy or sad) by button-press, using MEG compatible response pads. Variable intertrial intervals followed the presentation of the image.

**Figure 1.**
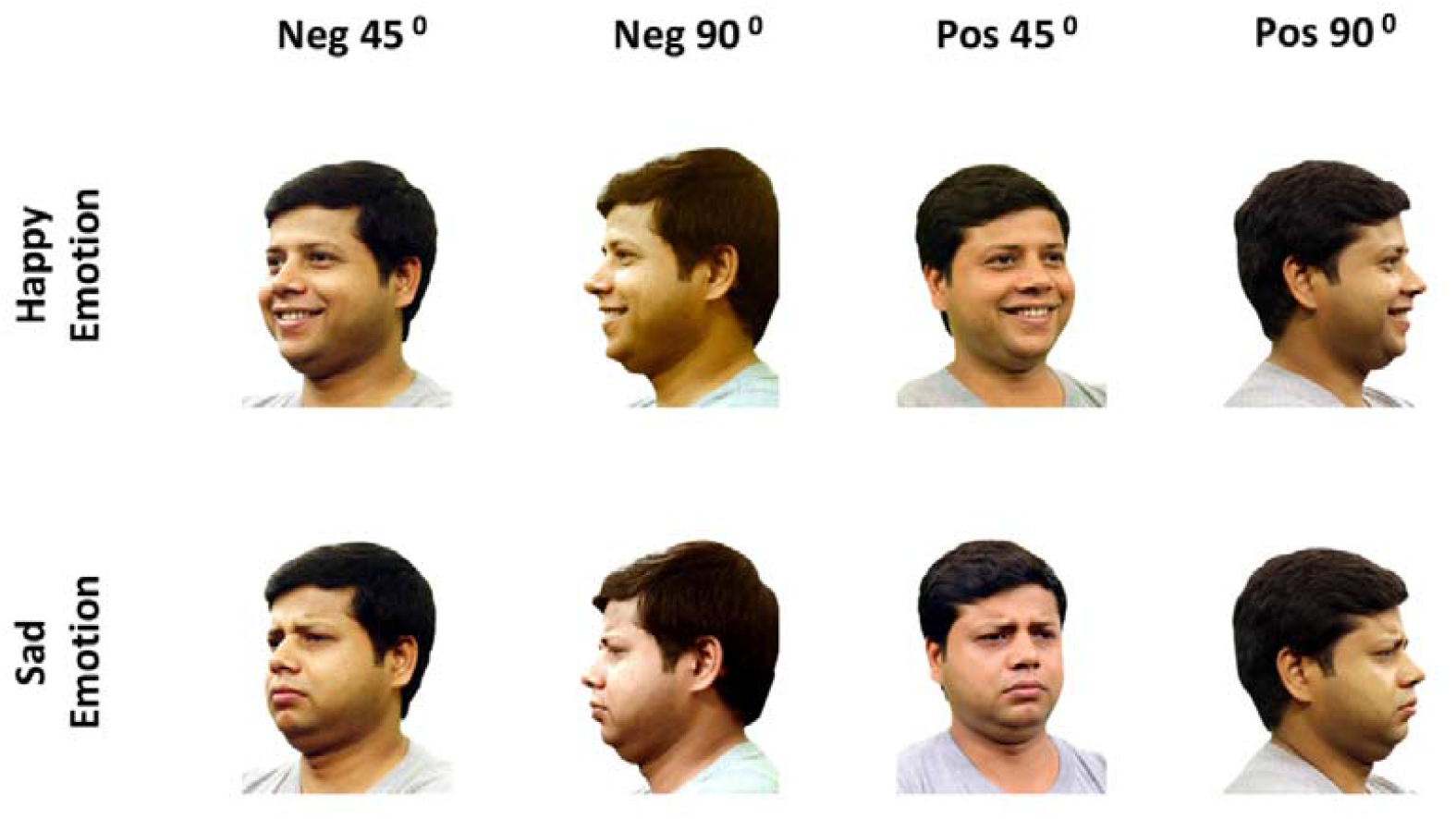
Faces at four different angles; Negative Visual angles i.e., Left Side of human face (−45^0^, -90^0^) and Positive Visual angles i.e., Right Side of human face (+45, +90^0^) in two emotions (Happy and Sad).

### Data acquisition and preprocessing

MEG data were acquired in a magnetically shielded room using the whole-scalp MEG system (Elekta-Neuromag, Helsinki, Finland) installed at the Intelligent Systems Research Centre (ISRC). The system is equipped with 102 sensor triplets (each comprising a magnetometer and two orthogonal planar gradiometers) uniformly distributed around the head of the participant. Head position inside the helmet was continuously monitored using four Head Position Indicator (HPI) coils. The location of each coil relative to the anatomical fiducials (nasion, left and right preauricular points) was defined with a 3D digitizer (Fastrak Polhemus, Colchester, VA, USA). This procedure is critical for head movement compensation during the data recording session. Digitization of the fiducials plus ∼300 additional points evenly distributed over the scalp of the participant which will be used during subsequent data analysis to spatially align the MEG sensor coordinates with T1 magnetic resonance brain images. MEG recordings were acquired continuously with a bandpass filter at 0.01-330 Hz and a sampling rate of 1 kHz. Eye movements were monitored with two pairs of electrodes in a bipolar montage placed on the external sides of each eye (horizontal electrooculography (EOG)) and above and below the right eye (vertical EOG). Similarly, cardiac rhythm was monitored using two electrodes, placed on the right side of the participants’ abdomen and below the left clavicle. Continuous MEG data were pre-processed off-line using the temporal Signal-Space-Separation (t-SSS) method (Taulu and Simola, 2006) which suppresses external interferences. MEG data were also corrected for head movements, and bad channel time courses were reconstructed using interpolation algorithms implemented in the software. Subsequent analyses were performed using Matlab R2014b (Mathworks, Natick, MA, USA).

### Sensor level ERF

MEG trials were corrected for jump artifacts and muscle artifacts. Heartbeat and EOG artifacts were detected using Independent Component Analysis (ICA) and linearly subtracted from recordings. The ICA decomposition was performed using the Fastica algorithm implemented in the Fieldtrip toolbox (Oostenveld et al., 2011), thirty (30) ICA components per subject were computed. ICA components having maximum correlation with EOG and EKG were automatically removed. On average 2 components were removed per participant. The artifact-free data were bandpass filtered between 0.5 and 45 Hz. Trials were segmented starting from 200 ms before the image presentation until 1000 ms after the onset of image presentation. Event Related Fields (ERFs) were quantified as the absolute amplitude of the 102 orthogonal planar gradiometer pairs by computing the square root of the sum of squares of the amplitudes of the two gradiometers in each pair. Baseline correction was also applied to the evoked data using a 200 ms window prior to presentation of an image at the beginning of each trial. The trial combination was performed in a 2 * 2 design (Emotions * Visual angles). Table 1 shows the contrast generated for each trial type.

**Table 1.** Trial combination and comparisons

|  | **Positive Visual Angle /**<br>**Right Side of Human**<br>**Face** | **Negative Visual Angle**<br>**/ Left Side of Human**<br>**Face** | **Contrast** |
| --- | --- | --- | --- |
| **Happy** | Happy Emotion<br>positive visual angles | Happy Emotion<br>negative visual angles | Positive vs<br>Negative visual<br>angles presenting<br>Happy Emotion |
| **Sad** | Sad Emotion positive<br>visual angles | Sad Emotion negative<br>visual angles | Positive vs<br>Negative visual<br>angles presenting<br>Sad Emotion |
| **Contrast** | Happy vs Sad Emotion<br>presented at Positive<br>visual angles | Happy vs Sad Emotion<br>presented at Neg visual<br>angles |  |

In the 2*2 design in Table 1, the comparison across columns illustrates the contrast in Emotions, whereas comparison across rows illustrates the contrast in visual angles. For the sensor level statistical analysis, a cluster-based permutation test was applied to evoked data corresponding to different contrasts explained in table 1. Four different comparisons were carried out. In the first comparison, we contrasted the ERFs for the Happy and Sad Emotions presented at positive visual angles. In the second comparison, we compared the ERFs for the Happy and Sad Emotions presented at Negative Visual angles. These two comparisons were made to determine differences in emotion processing in response to either the right (Positive Visual angles) or the left (Negative Visual angles) side of the human face. Then, we compared the ERFs for the Positive and Negative Visual angles presenting Happy Emotions. The last comparison contrasted the ERFs of Positive and negative visual angles presenting Sad Emotions. These comparisons were used to assess how well the information about each emotion was processed depending on which side of the face the expression of the emotion was viewed. In all cases, differences between conditions were analyzed using a clustering and randomization test (Maris and Oostenveld, 2007). A randomization distribution of cluster statistics was constructed for each subject over time and sensors and used to evaluate whether there were statistically significant differences between conditions over participants. In particular, t-statistics were computed for each sensor (combined gradiometers) and time point during the 0 - 700 ms time window, and a clustering algorithm formed groups of channels over time points based on these tests. The neighborhood definition was based on the template for combined gradiometers of the Neuromag-306 provided by the toolbox. In order for a data point to become part of a cluster, a threshold of *p* = 0.05 was applied (based on a two-tailed dependent t-test, using probability correction) and it had to have at least two neighbours. The sum of the t-statistics in a sensor group was then used as a cluster-level statistic (e.g., the maxsum option in Fieldtrip), which was then tested using a randomization test using 1000 runs.

### Source Reconstruction

Source reconstruction mainly focused on the statistically significant effects observed at the sensor-level in the time range 370 – 420 ms after presentation of the stimuli. MEG-MRI co-registration was performed using interactive closet point algorithm implemented in fieldtrip software (Oostenveld et al., 2011). T1-weighted MRI SPM template was segmented into scalp, skull, and brain components using the SPM segmentation algorithms implemented in fieldtrip. The source space was defined as a regular 3D grid with a 5 mm resolution and the lead fields were computed using a single-sphere model for 3 orthogonal source orientations. The lead field was also normalized to reduce the center-head bias issue in beamformers and warped to MNI source space. Whole brain source activity was estimated using a linearly constrained minimum variance (LCMV) beamformer approach (Veen et al., 1997). Only planar gradiometers were used for inverse modelling. The covariance matrix used to derive LCMV beamformer (Common filter) weights was estimated from the pre- and post-stimulus data in the pre-stimulus (from - 200 ms prior to presentation of emotional face stimuli) to post-stimulus (750 ms after the presentation of the emotional face stimuli) time range.

The LCMV beamformer focused on the (baseline corrected) evoked data in the time period 370 –420 ms post-stimulus (when the mean ERF peak amplitude difference across participants was statistically different between two conditions at the sensor level, and was largest). The group activity across participants was calculated by averaging the source map from all the participants. The neural activity index (NAI) was calculated as relative percent change in pre- and post-data ((Data _Post_ – Data _Pre)_. / Data _Pre)_ * 100) for the difference signal between Sad and Happy Emotion. The resulting NAI was interpolated with the AAL brain (Tzourio-Mazoyer et al., 2002) atlas and plotted on an inflated brain. To avoid “double dipping”, we did not compare the source maps for different conditions.

### Multivariate pattern analysis (MVPA)

Time-resolved within-subjects multivariate pattern analysis was performed to decode mainly two features i.e., the emotion in the presented stimuli (Happy or Sad) and the visual angle (Positive or Negative), from the MEG data. This within-subject classification has an advantage over other methods; the classification algorithm may leverage individual subject specific characteristics in neural patterns since the classifiers do not need to generalize across different subjects. The data were segmented from 100 ms prior to the onset of the stimuli to 850 ms after the stimuli presentation. The data were classified separately for both the features using a linear support vector machine (SVM) classifier with L2 regularization and a box constraint of c=1. The classifiers were implemented in Matlab using the LibLinear package (Fan et al., 2008) and the Statistics and Machine Learning Toolbox (Mathworks, Inc.). We performed a binary classification of the Face Emotions. The same process was repeated for decoding the Visual angles from the MEG data. The data were down sampled to 200 Hz prior to the classification. Pseudo trials were generated to improve the signal-to-noise-ratio (SNR) by creating trials’ bins, resulting in a set of 10 trials for each bin (Dima and Singh, 2018; Nara et al., 2021). This pseudo trial generation was repeated 100 times to generate trials with a higher signal to noise ratio. The data were then randomly partitioned using 5-fold cross-validation. The classifier was trained on 4-folds and tested on 1-fold and this process was repeated until each fold is left out once. The procedure of generating pseudo trials, dividing the data into 5 folds, and training and testing classifiers at every time point was repeated 25 times; classification accuracies were then averaged over all these instances to yield more stable estimates. To improve data quality, we also performed multivariate noise normalization (Guggenmos et al., 2018). The time-resolved error covariance between sensors was calculated based on the covariance matrix of the training set and used to normalize both the training and test sets in order to down weight MEG channels with higher noise levels. Cluster corrected sign permutation tests were applied to the accuracy values obtained from the classifier with cluster-defining threshold p < 0.05, corrected significance level i.e., cluster-alpha p < 0.01. We used t-tests to evaluate differences between conditions across participants.

## Results

### Evoked analysis Results

First, we compared the Happy and Sad emotions at Visual angles. This analysis found a significant negative cluster (*p* = 0.006) in time range 372 – 419 ms. There were two other clusters which were marginally significant, the first (*p* = 0.063) emerges from 303 – 341 ms, and the second (*p* = 0.079) emerges from 437 – 466 ms. The neural responses to sad emotion were higher in amplitude compared to the happy emotion (Figure 3A). The cluster was located in fronto-temporal brain areas (Figure 3B) and reflects the N400 effect. This effect was not significant when happy and sad emotions were compared at negative visual angles, suggesting that the happy and sad emotions were differentiated from Positive Visual angles only. The third comparison contrasted the neural responses to Positive and Negative Visual angles during processing of happy emotion; however, this comparison did not yield a statistically significant effect. The fourth comparison compared the neural responses to Positive and Negative visual angles during processing of sad emotion, this comparison was also not statistically significant.

**Figure 2.**
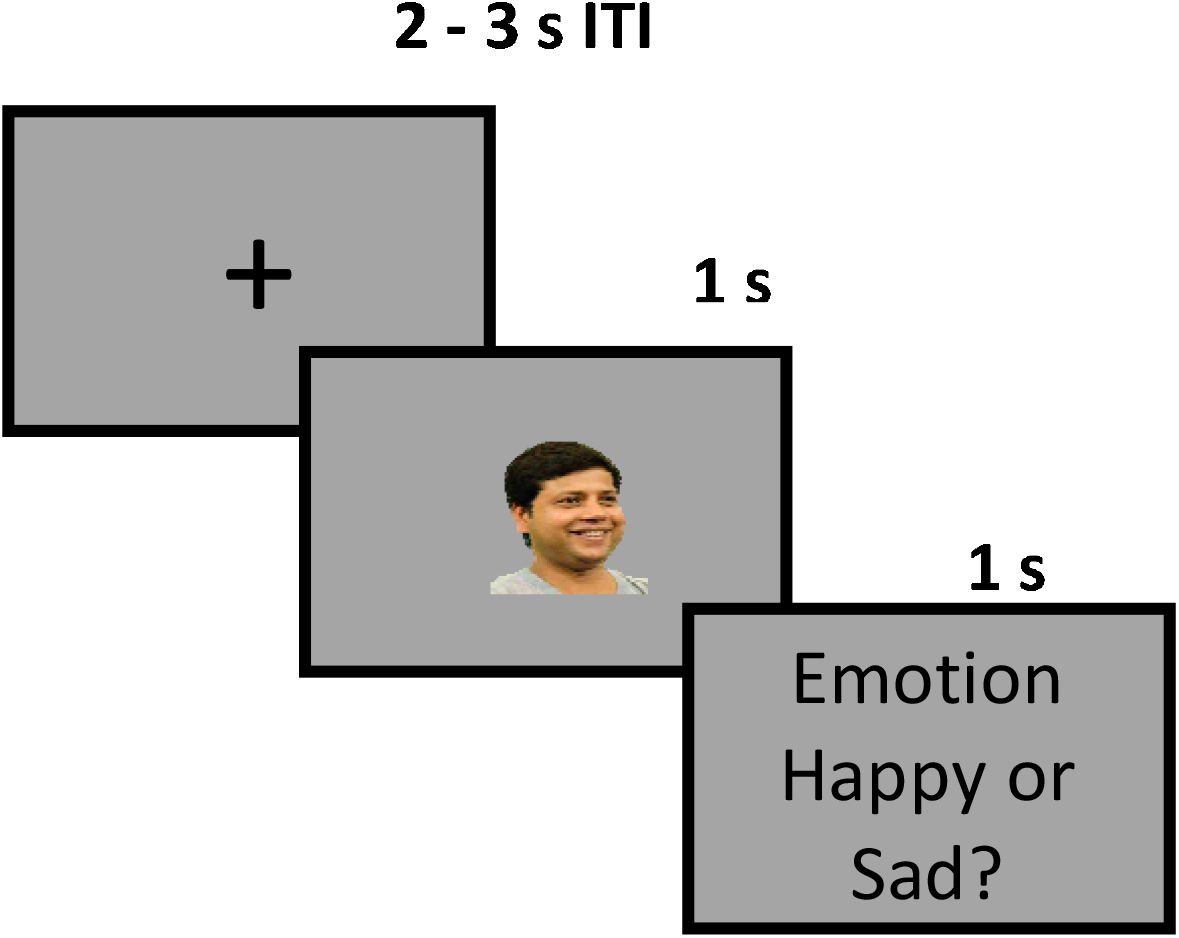
The Experimental Design of single trial is shown here.

**Figure 3.**
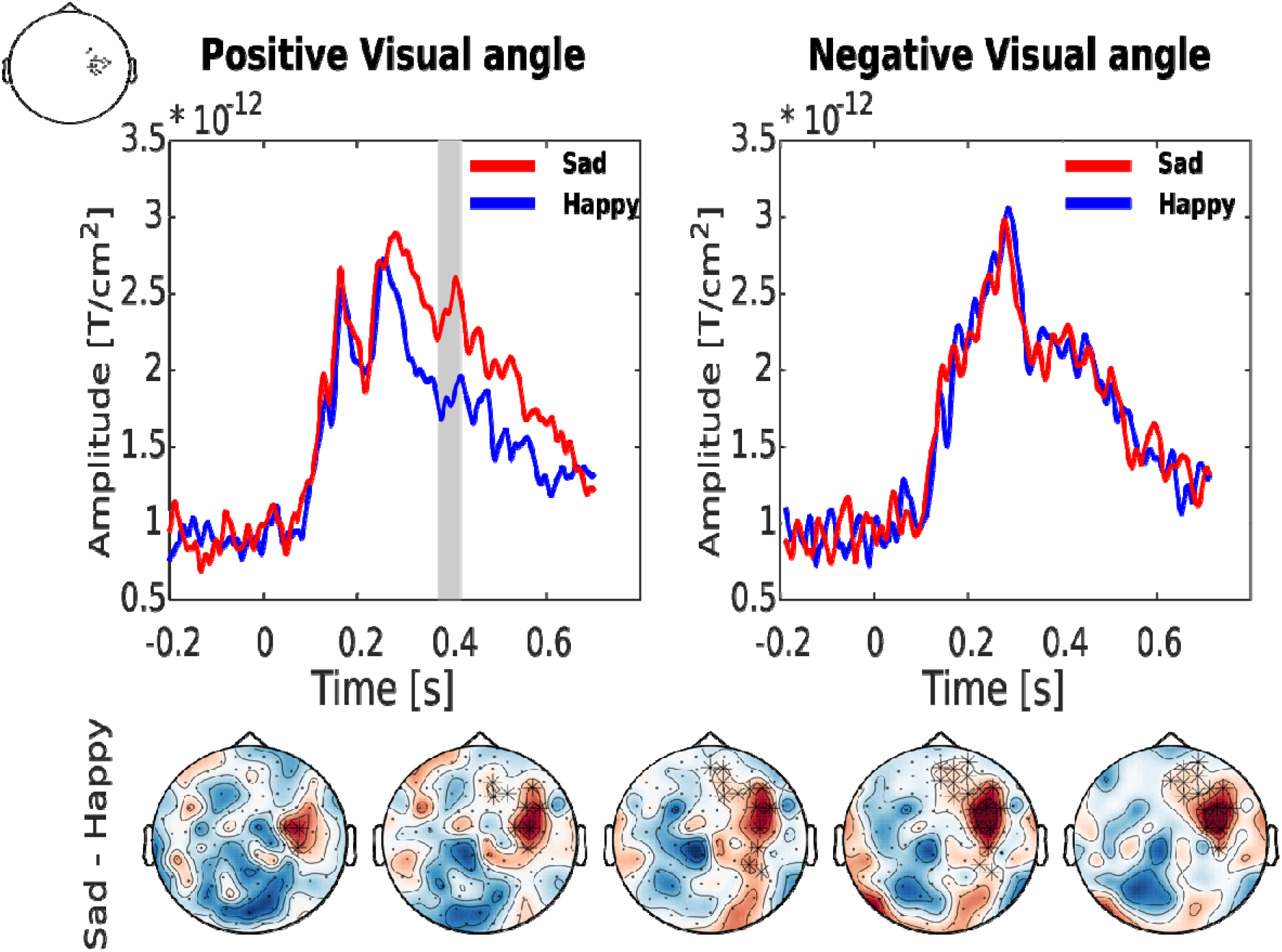
(Top) Sensor level ERF [averaged from 6 sensors: ‘MEG1122+1123’, ‘MEG1132+1133’, ‘MEG1332+1333’, ‘MEG1342+1343’, ‘MEG2222+2223’, ‘MEG2412+2413’] for the Happy and Sad Emotion measured at Positive and Negative Visual angles. The grey box shows the time points where the amplitude of ERFs (in time window 0-750 ms) was higher (*p* < 0.05, cluster-based permutation test) for Sad Emotion compared to the Happy emotion. (Bottom) The topoplot shows the difference of ERFs in Sad and Happy emotion. The asterisks (*) shows the statistically significant channel emerging from cluster-based permutation test (time window = 370–420 ms, post stimulus onset).

### Source reconstruction

We next identified the brain regions underlying the relevant effects observed at the sensor level. Source activity was estimated from sensor-level ERFs in the 370–420 ms interval. Whole-brain maps of source activity were created for differences in neural activity (emerging from Sad – Happy). Figure 4 shows the source activity localized in both ventral and dorsal stream of visual processing. The brain areas showing maximum NAI differences include Fusiform gyrus, lingual gyrus, putamen and Pre and Post central gyrus. These brain areas have been proposed to be involved in face and emotion recognition. This suggests that the Positive Visual angles were able to activate the brain areas relevant to face and emotion recognition.

**Figure 4.**
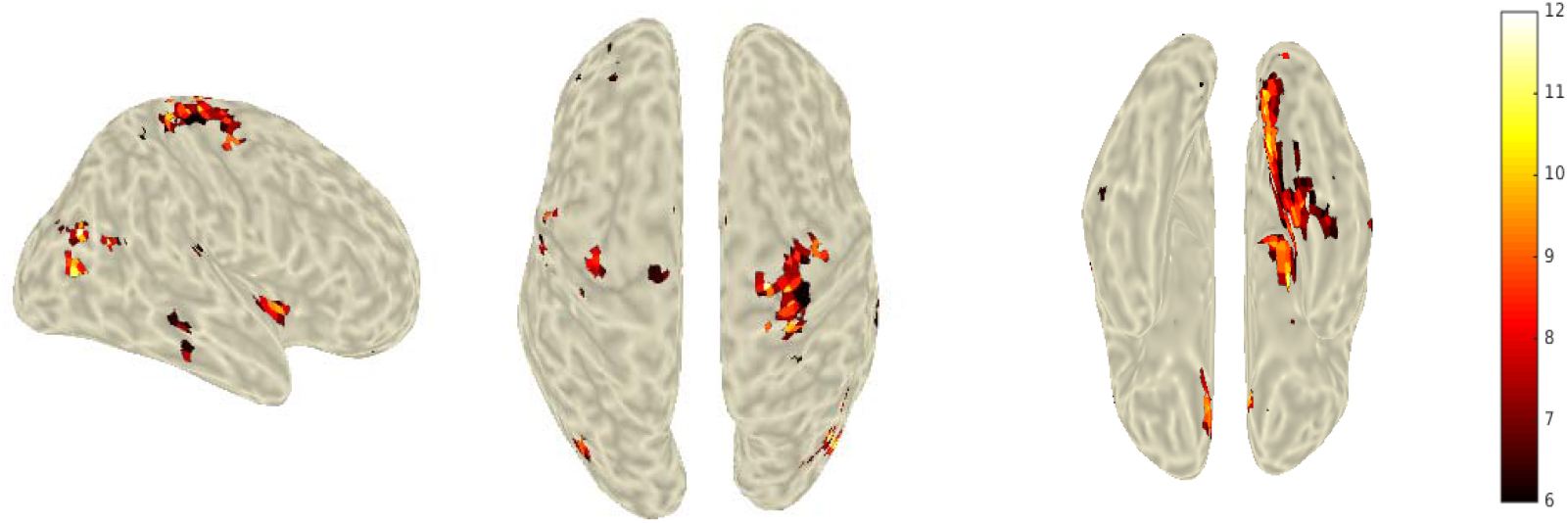
Source reconstructed brain maps showing the Neural activity Index (NAI).

### MVPA Results

The first comparison was performed to investigate the temporal dynamics of emotion processing at both Positive and Negative Visual angles. MVPA revealed that Happy emotions are processed faster (facilitating the earlier decoding of visual angles i.e., Positive vs Negative Visual angle) compared to the Sad emotions. Figure 5 (left) shows that the decodable information relating to the Happy emotion began to emerge as early as 70 ms, whereas the same information for visual angles presenting Sad emotion did not begin to emerge until 100 ms post stimulus-onset. The decoding accuracy for the Happy emotion was also higher than the Sad emotion (*p*= 0.031, one tailed *t*-test). The second comparison was carried out to investigate the temporal dynamics of emotion processing (i.e., Happy or Sad) imposed by Positive or Negative visual angles. Figure 5 (right) shows that the information about the content of the emotion is only disclosed when viewed from Positive Visual angles. This information is revealed from 420 – 450 ms after the presentation of the stimuli. The decoding accuracy was higher and statistically significant for Positive Visual angles compared to Negative visual angle (*p* = 0.0019, one tailed *t*-test).

**Figure 5.**
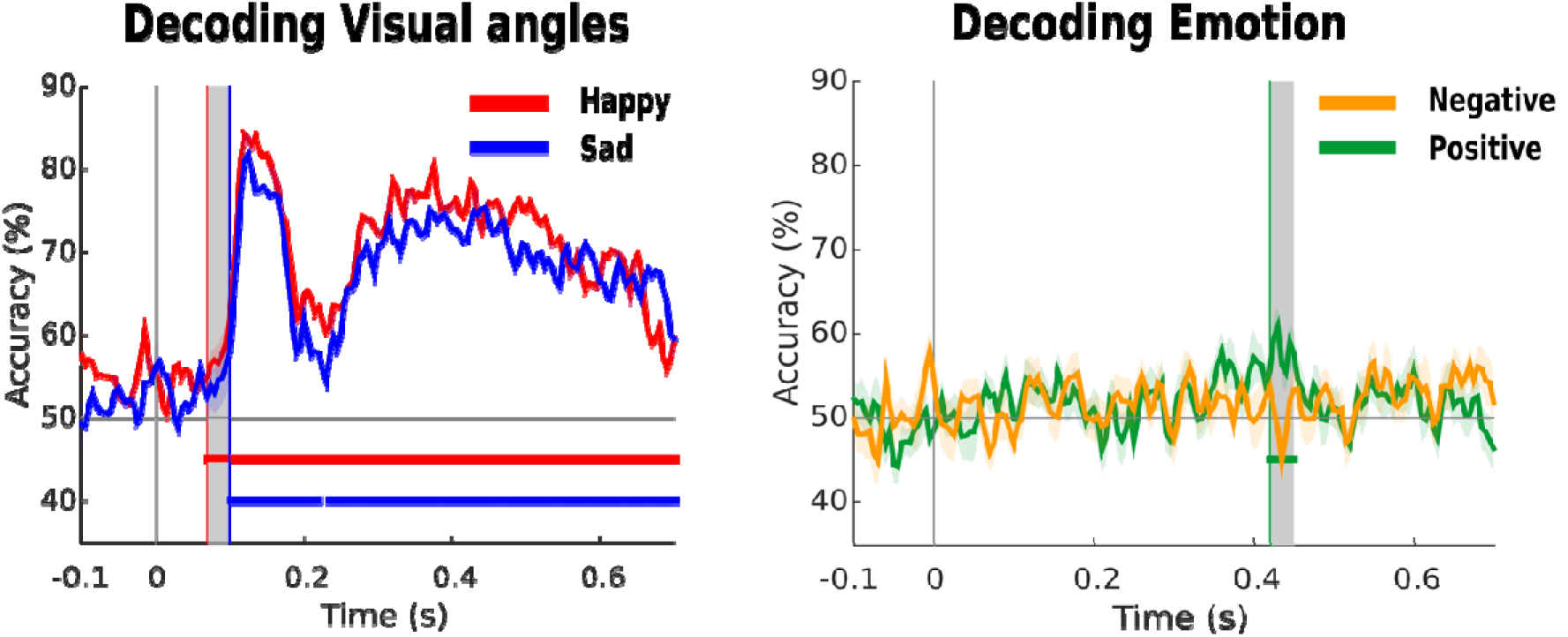
(Left) Time resolved decoding accuracy for visual angle (classifier trained to discriminate between Positive and Negative visual angles) used to present both Happy and Sad emotion. The dotted red and blue line show the statistical significance (p < 0.05, cluster-based permutation test). The grey shaded region shows statistically significant time duration when decoding accuracy was higher for the Happy condition compared to the Sad condition. B) Time resolved decoding accuracy for emotion processing (classifier trained to discriminate between Happy and Sad emotion) when presented at either Positive or Negative visual angles. The dotted green line shows the statistical significance (p < 0.05, cluster-based permutation test). The grey shaded region shows the statistically significant time window during which decoding accuracy was higher for Positive Visual angle presentations compared to Negative.

## Discussion

To the best of our knowledge, this is the first study investigating different facial angles and emotions together, using MEG. The findings clearly show neurophysiological evidence for hemispheric and hemifacial asymmetry in the processing of facial expressions of emotions. The other two studies using the same database presented only the findings of behavioral data (Sharma and Bhushan, 2019) or mix of the behavioral data and geometric analysis of these expressions (Bhushan and Munshi, 2021). Both of them could not examine hemispheric or hemifacial asymmetry, although they had a unique database with facial expressions captured from multiple angles.

This study reveals two important findings. First, the right side of the human face (positive visual angles) provides more information about the emotional content – as illustrated in the topoplots presented in Figure 3, which show differential activation of the right hemisphere. Second, the brain processes the information about the facial angles (Happy and Sad emotions) in different time frames. Happy expressions are processed faster compared to sad expressions. This is supported by the N400 effect observed in the ERF analysis and the absence of significant decoding latency in the MVPA analysis. Both the studies (Sharma and Bhushan, 2019; Bhushan and Munshi, 2021) have reported higher accuracy rate for recognition of happiness, but this study confirms difference in terms of processing time as well.

Further, in sharp contrast to the findings of earlier behavioral studies (Mandal et al., 2001; Bhushan, 2006) supporting hemifacial bias, which found the left side of the face to distinctly express negative emotions, our findings suggest that the right side of the face provides distinct information for perception of emotion, while activation is increased in the right hemisphere during this process.

These findings both agree and disagree with the right-hemisphere hypothesis – support for which has found the right hemisphere is dominant for negative emotions whereas the left hemisphere is dominant for positive emotions (Silberman and Weingartner, 1986; Heilman and Bowers, 1990; Yecker et al., 1999). The current findings relating to hemispheric dominance are better aligned with previous research on the perception of emotion, which has also evidenced right hemisphere dominance irrespective of emotional valence (Bryden M. P., 1982; Davidson, 1984). Although Bryden and Davidson reported the findings on the basis of the study of the frontal view of the face, here the effect has been found to hold for the positive angular profile (although not for negative visual angle presentation).

Mandal and Ambady (Mandal and Ambady, 2004) have argued that the left side of the face displays culture-specific signals, whereas the right side of the face displays universal emotional signals. As mentioned earlier, one of the major challenges regarding facial expression databases of basic emotions is the lack of a culture-fair database. This would demand universal emotional signals on the facial expression. Our findings suggest that the happy and sad facial expressions of IAPD exhibit universal emotional signals and thus are suitable for pan-cultural administration.

Another important finding of this study is the faster processing of happy emotion (facilitating the earlier decoding of visual angles, i.e., positive vs negative visual angle) compared to sad emotion. An earlier study using the IAPD (Sharma and Bhushan, 2019), found happy emotion to be the “easily identifiable emotion”. However, this was a behavioral study which reported a hit rate of 99.91 (SD 0.277) for happy and 94.18 (SD 2.99) for sad expressions. It reported comparable unbiased hit rate (Hu Happy= .88, Hu Sad= .8). While adding fractal geometry to IAPD database, Bhushan and Munshi (Bhushan and Munshi, 2021) analyzed if the geometric changes in the images depicting facial expressions affect the recognition accuracy of these emotions or not. They found happy expression to be isotropic, i.e., recognition of happy emotion does not get affected by visual angle. The present study suggest that recognition of happy expressions is affected by the positive visual angle that too in a comparatively shorter timeframe.

The novelty of this study lies in the finding that the time frame needed for the processing of happy emotions is shorter than that for sad expressions of emotion. The decodable information for happy emotion was found to emerge much earlier (70 ms) compared to sad emotion (100 ms). For the negative cluster the time ranged between 372 – 419 ms, with two other clusters (303 – 341 ms and 437 – 466 ms) observed before and after the significant cluster – thus, appearing to be part of the same cluster emerging from the N400 effect. The neural responses to the sad emotion images elicited a higher amplitude ERF, compared to the happy emotion in the fronto-temporal area, reflecting the N400 effect.

The N400 effect is mainly reported in linguistic experiments where this component emerges if the word has an emotional content. The N400 effect we observe here also arises from the emotional content of the stimuli, however there is the possibility that the source generators of this effect lie in the temporal area.

Earlier studies have shown right hemisphere dominance for perception of facial expressions of emotion especially basic emotions(Alfano and Cimino, 2008; Vytal and Hamann, 2010) found activation of superior frontal gyrus while processing sadness and the right superior temporal gyrus while processing happiness. Fusiform gyri activation for static images has been reported by other researchers as well (Vandewouw et al., 2020). One of the novel findings of this study is finding the shorter time frame needed for decoding the happy expressions vis-à-vis the sites responsible for it. The NAI differences were found in the fusiform gyrus, lingual gyrus, putamen and Pre and Post central gyrus suggesting the involvement of these areas in the recognition of facial expression of emotions, specifically by the facial stimuli depicting positive visual angles.

### Limitations

Although the experimental design, data collection, and data analysis were carried out very carefully, as this is a novel investigation – replication of results will be required for further validation. The authors do accept the limitations of the number of participants (thirteen here), unfortunately data collection was cut short due to Covid 19.

## Conclusion

The brain takes shorter time to process happy expressions compared to sadness, specifically the facial stimuli depicting positive visual angles only. It activates fusiform gyrus, lingual gyrus, putamen, and pre- and post-central gyrus.

## Acknowledgement

This research was supported by the project BES-2016-077560 funded by the Spanish Ministry of Economy and Competitiveness (MINECO) awarded to SN and NM. SN does acknowledge the support from EMBO short term fellowship. BB was partly supported through Science & Engineering Research Board (SERB), Govt. of India, grant no.

MTR/2019/000224. The authors thank Sujit Roy, the lab staff for helping in data acquisition and our participants for their valuable contribution to this study. SN does acknowledge the support from Dr Mikel Lizarazu and Dr Craig Richter for helping in source reconstruction method implemented in this study. This work was supported in part by Northern Ireland Functional Brain Mapping (NIFBM) Facility Project funded through InvestNI and the Ulster University under the Grant 1303/101154803 and the MRC UK MEG Partnership Grant MR/K005464/1.

## Data and Code Availability Statement

The MATLAB scripts used for analyzing the data and the full dataset is available upon requests directed to Dr. Girijesh Prasad or Sanjeev Nara

